# Why do pathway methods work better than they should?

**DOI:** 10.1101/2020.07.30.228296

**Authors:** Bence Szalai, Julio Saez-Rodriguez

## Abstract

Different pathway analysis methods are frequently applied to cancer gene expression data to identify dysregulated pathways. In most cases these methods infer pathway activity changes based on the gene expression of pathway members. However, pathways are constituted by signaling proteins, and their activity - not their abundance - defines the activity of the pathway; the association between gene expression and protein activity is in turn limited and not well characterised. Other methods infer pathway activity from the expression of the genes whose transcription is regulated by the pathway of interest, which seems a more adequate proxy of activity. Despite these potential limitations, membership based pathway methods are frequently used and often provide statistically significant results.

Here, we submit that pathway based methods are not effective because of the correlation between the gene expression of pathway members and the activity of the pathway, but because pathway member gene sets overlap with the genes regulated by transcription factors (regulons). This implies that pathway methods do not inform about the activity of the pathway of interest, but instead the downstream effects of changes in the activities of transcription factors.

To support our hypothesis, we show that the higher the overlap to transcription factor regulons, the higher the information value of pathway gene sets. Furthermore, removing these overlapping genes reduces the information content of pathway gene sets, but not vice versa. Our results suggest that results of classical pathway analysis methods should be interpreted with caution, and instead methods using pathway regulated genes for activity inference should be prioritised.

**Graphical abstract:** 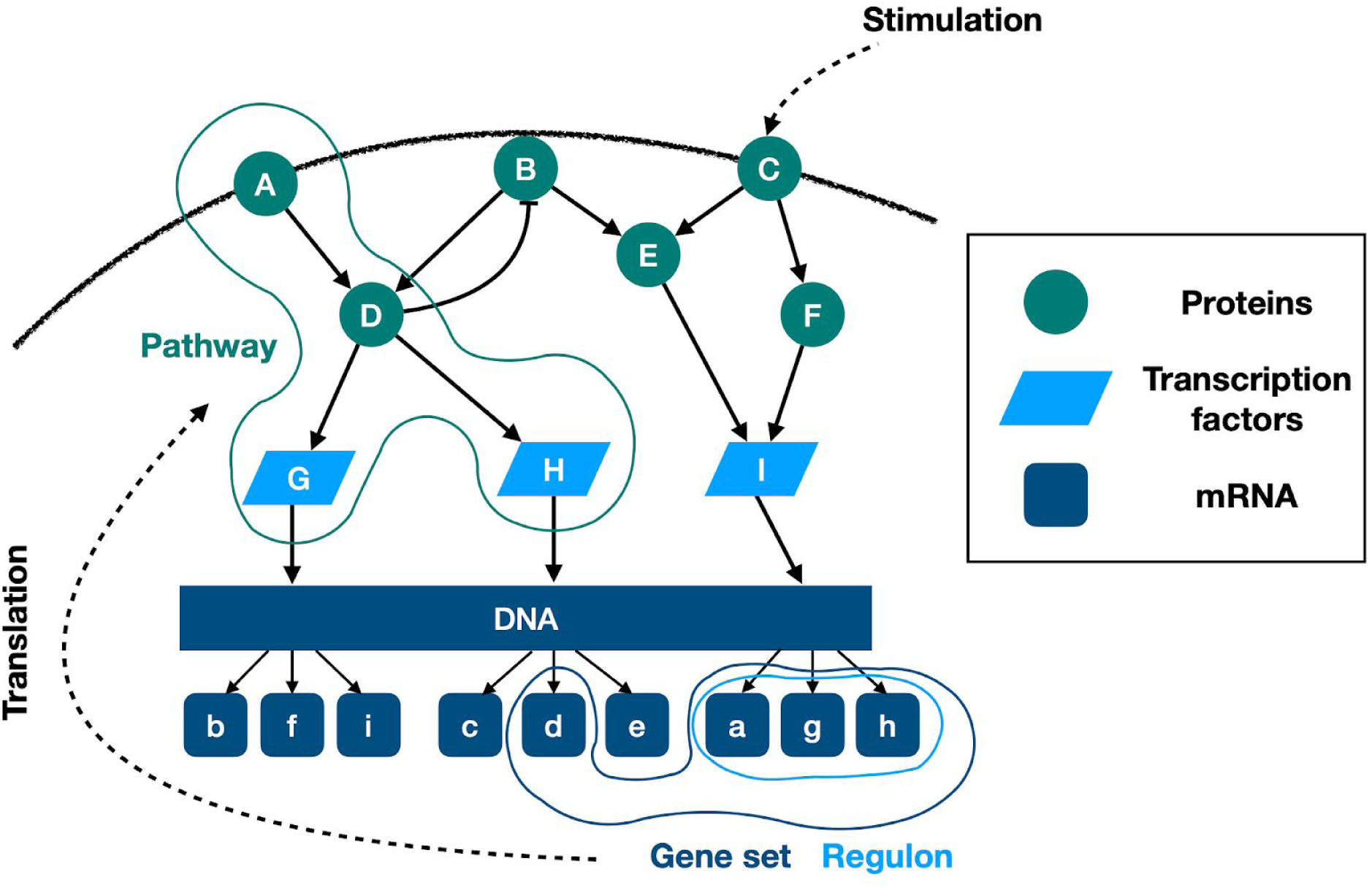

## 1. Introduction

Identification of driver events, such as mutations and copy number alterations in cancer genomes is crucial to understand mechanisms behind cancer initiation, and to identify new therapeutic strategies. However, cancer is not only a genetic disease, but also a disease of altered signaling (Yaffe 2019). Signaling pathway activity is mediated by protein - protein interactions and post-translational modifications (PTMs, e.g.: phosphorylation). Understanding pathway activity alterations in cancer is important to identify new personalised therapeutic options, especially as driver mutations are not always good biomarkers (Smith and Sheltzer 2018). Pathway activities also show comparable or even stronger associations with the drug sensitivity of cancer cell lines, than mutations (Schubert et al. 2018).

While the natural choice of data type for describing pathway activity is proteomics / phosphoproteomics (Saez-Rodriguez and Blüthgen 2020), due to the much higher abundance (and more affordable acquisition) of transcriptomics (gene expression) data, the latter one is also frequently used to infer pathway activities. In a typical scenario, differential expression is calculated between two groups of samples (e.g.: cancer vs. healthy tissue), followed by some kind of gene set enrichment (GSE) method to identify dysregulated groups of genes (Subramanian et al. 2005). If the gene set of interest consists of the genes of a signaling pathway, the result of GSE is typically interpreted as altered pathway activity. Several recent methods not only use the gene sets, but also the interaction between the (protein product) of these genes (i.e. network topology), to incorporate information flow (activatory and inhibitory interactions between the proteins) into the analysis (Hidalgo et al. 2017; Nguyen et al. 2019).

It is important to highlight that using classical pathway analysis methods with gene expression data (directly or indirectly) assumes a strong correlation between gene expression, protein abundance and protein activity. Pathway gene sets are defined based on groups of protein products, while transcriptomics measures mRNA abundance. The exact correlation between gene expression and protein levels depends on multiple biological and technical factors and remains an area of investigation, but is clearly limited (Y. Liu, Beyer, and Aebersold 2016; Buccitelli and Selbach 2020). For example, across a large panel (375) of cancer cell lines (Nusinow et al. 2020), the average (Pearson’s) correlation (between 12,755 gene expressions and protein abundances) does not exceed 0.5. Connecting gene expression to protein activity is even more complicated, as protein activity is controlled by factors ranging from localization to post-translational modifications (PTMs), and gene expression can be even incoherent with the expected flow of information in signaling networks (Larsen et al. 2019; Piran et al. 2020). Accordingly, predictability of proteomic and phosphoproteomic levels from genomics and transcriptomics is rather modest (Yang et al. 2020).

In contrast to these pathway “mapping” based methods, “footprint” based pathway analysis techniques do not infer pathway activity from the gene expression of pathway members, but from expression of genes regulated by the pathway of interest (Dugourd and Saez-Rodriguez 2019). These methods have shown increased performance compared to classical pathway analysis techniques in several benchmarks (Parikh et al. 2010; Schubert et al. 2018; Holland et al. 2020). A recent systematic analysis of gene sets (Cantini et al. 2018) also showed that footprint based gene sets have higher information value than pathway mapping based gene sets. In this study “informative gene sets” were defined by their ability to rank samples based on some biological phenotype (e.g.: cancer vs. healthy samples).

Despite these theoretical considerations, classical pathway analysis methods with gene expression data are one of the most frequently used computational methods in systems biology, and they frequently produce significant results - for example, pathway gene sets are often found to be significantly more altered in gene expression data than randomly expected. In this manuscript we propose a possible mechanism for this effectiveness of mapping based pathway analysis methods.

We propose that pathway gene sets are not informative because they infer the activity of the corresponding pathway, but because they are overlapping with the regulated genes of transcription factors (TFs) (Figure 1). TF activity can be effectively inferred from gene expression data, using the expression of the regulated genes of the transcription factors (regulon) (Garcia-Alonso et al. 2018; Alvarez et al. 2016). Expression of TF regulons can thus be viewed as footprints of TF activity. TF regulons are informative gene sets, as shown also in several benchmarks (Garcia-Alonso et al. 2019; Holland, Szalai, and Saez-Rodriguez 2019; Keenan et al. 2019; Holland et al. 2020). We reasoned that if a pathway gene set has significant overlap with a TF regulon (i.e. the TF regulates the expression of pathway members), the pathway gene set will be also informative. However, in this case, the gene set enrichment score does not reflect the activity of the signaling pathway, but reflects the corresponding transcription factor activity. To investigate this hypothesis, we calculated similarity between TF regulons (from DoRothEA resource (Garcia-Alonso et al. 2019)) and pathway mapping based gene sets. We also calculated the ability to identify biological meaningful subgroups in gene expression data of different gene sets, that we define as their information value. We show that informative pathway gene sets have higher similarity (overlap) to transcription factor regulons, while the reverse is not true - TF regulons can be informative without being similar to pathway gene sets. We also show that removing the genes that overlap with TF regulons from pathway gene sets decreases the informative value of pathway gene sets, while this has little effect on the informative values of TF regulons. Based on these results we submit that footprint based methods are a more adequate choice to infer pathway and TF activities from gene expression data, and that the effectiveness of pathway gene sets are mainly driven by their similarity to TF regulons.

**Figure 1.**
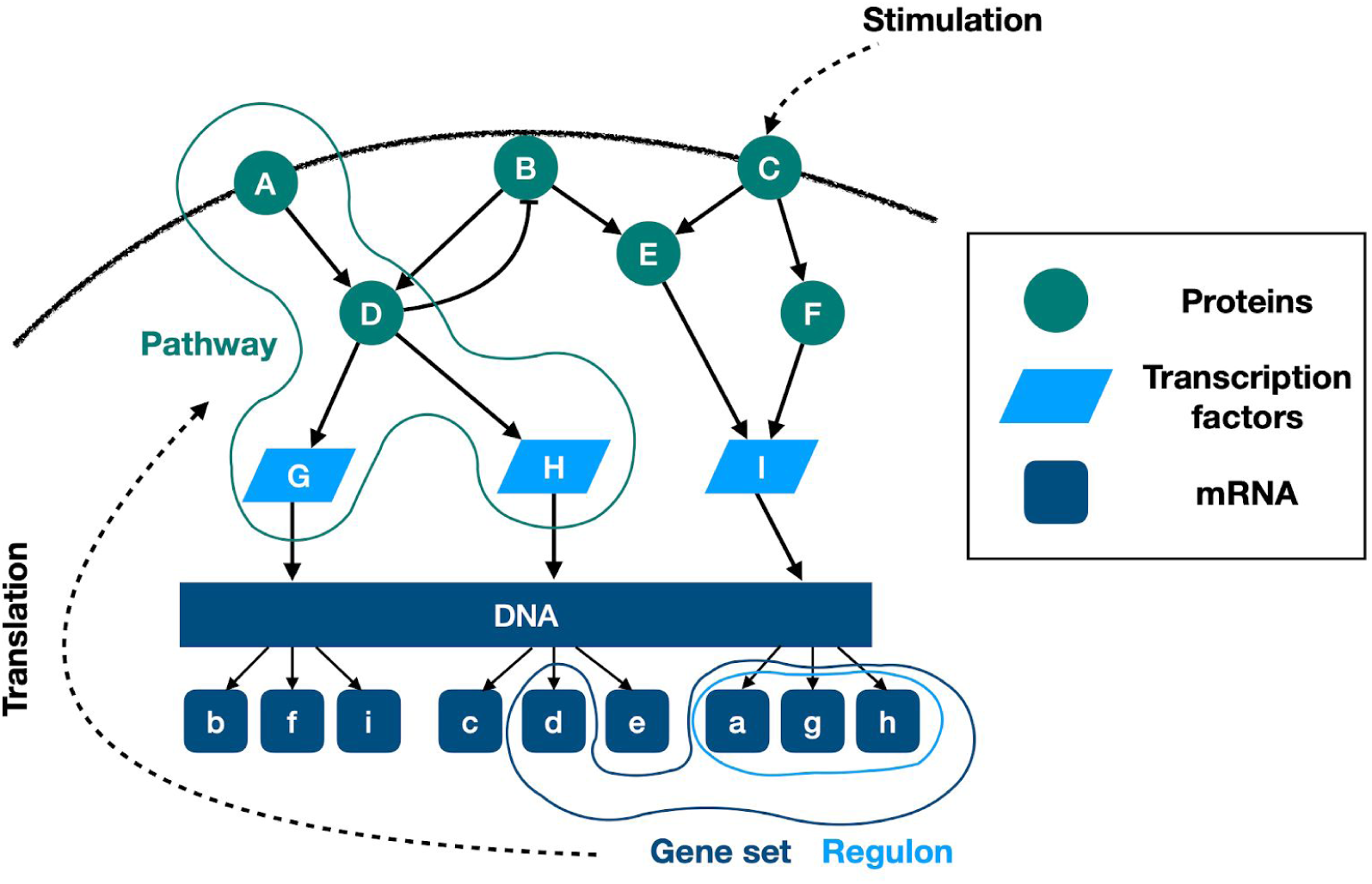
Concept of overlapping pathway gene sets and transcription factor regulons signaling networks (pathways) lead to transcription changes via the altered activation of transcription factors. Expressed mRNAs are translated to proteins. Activity of signaling can also be altered by other factors like perturbations, mutations etc. Pathways are defined on the level of interacting proteins. Regulons are the regulated gene of a transcription factor. Pathway mapping based gene sets (created by pathway member genes) are defined of the level of gene expression, and potentially overlap with transcription factor regulons.

## 2. Results

### 2.1 Pathway gene sets overlap with transcription factor regulons

We used three pathway mapping based gene set databases (KEGG (Kanehisa et al. 2019), Reactome (Jassal et al. 2020) and Biocarta, all downloaded from MSigDB (Subramanian et al. 2005)), one transcription factor regulon database (DoRothEA (Garcia-Alonso et al. 2019)) and one footprint gene set database (CGP, chemical and genetic perturbations gene sets from MSigDB) in this study. From DoRothEA we used high confidence (DoRothEA AB) and low confidence (DoRothEA CD) regulons. As some gene set databases had very large gene sets (SFigure 1), we focused only on gene sets with <= 250 genes, to make the different databases more comparable. We calculated similarity (Jaccard index or overlap coefficient, Methods) between all gene sets of selected database pairs (e.g. DoRothEA and Biocarta in Figure 2A), and calculated the maximum value of similarity for each gene set. We used maximal similarity, as we were interested in the similarity of a gene set to any other gene set of the other database. High confidence (AB) DoRothEA had higher maximal Jaccard index with the other gene set databases than randomly expected (Figure 2B, Mann-Whitney U p-values: 8.76e-26, 9.10e-17, 2.42e-21 and 3.75e-30 for KEGG, Biocarta, Reactome and CGP, respectively). We saw similar results using overlap coefficient, while low confidence DoRothEA had generally lower, but still significant difference compared to random gene sets (SFigure 2 and STable 1).

**Figure 2.**
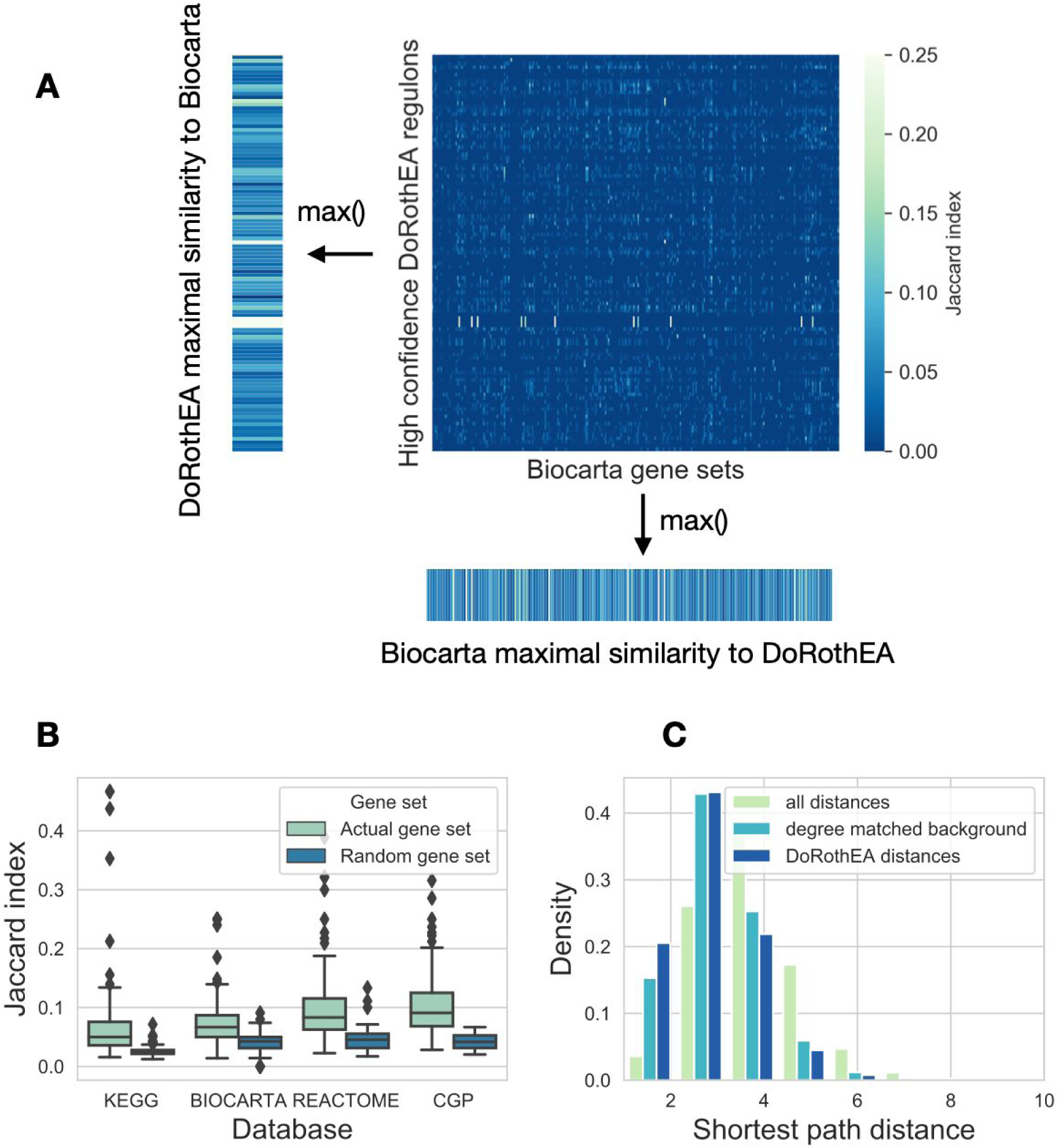
Similarity of gene sets. (A) Similarity (Jaccard index) matrix between gene sets of DoRothEA and Biocarta. For each row (DoRothEA gene sets) and column (Biocarta gene sets) the maximal similarity was calculated. (B) Distribution of maximal similarity values (Jaccard index, y axis) to high confidence DoRothEA regulons for the investigated gene set databases (x axis). Random gene sets (color code) were created by sampling gene sets (with corresponding size) from the genes of the given database. (C) Distribution of the shortest path distances between proteins transcriptionally regulated by the same transcription factor. For comparison the whole distribution of shortest path distances and degree matched background distribution is shown (color code).

As CGP collects perturbation related gene expression signatures, the high similarity of CGP and DoRothEA gene sets was not surprising. If a perturbation leads to activation / inhibition of a transcription factor, gene expression changes are expected to overlap with the regulon of the corresponding TF. However, as pathway gene sets are defined based on the protein-protein interactions of the products of the gene (not based on interactions among the genes driven by regulatory networks), their similarity to DoRothEA gene set was not self-evident.

Based on this similarity, we wondered whether protein products of genes regulated by a given TF also generally interact with each other in the (protein) signaling network. We calculated shortest path distances for the members of different gene sets, using the undirected OmniPath signaling network (Türei, Korcsmáros, and Saez-Rodriguez 2016). High confidence (AB) DoRothEA gene sets had lower intra gene set distance than both background and degree matched random distribution (Figure 2C, Mann-Whitney p values <1e-100). We observed similar results for pathway mapping based gene sets (SFigure 3), and a less pronounced effect for low confidence (CD) DoRothEA and CGP. In summary, our analysis of the gene set compositions showed that TF regulon gene sets have higher similarity to pathway based gene sets than randomly expected.

**Figure 3.**
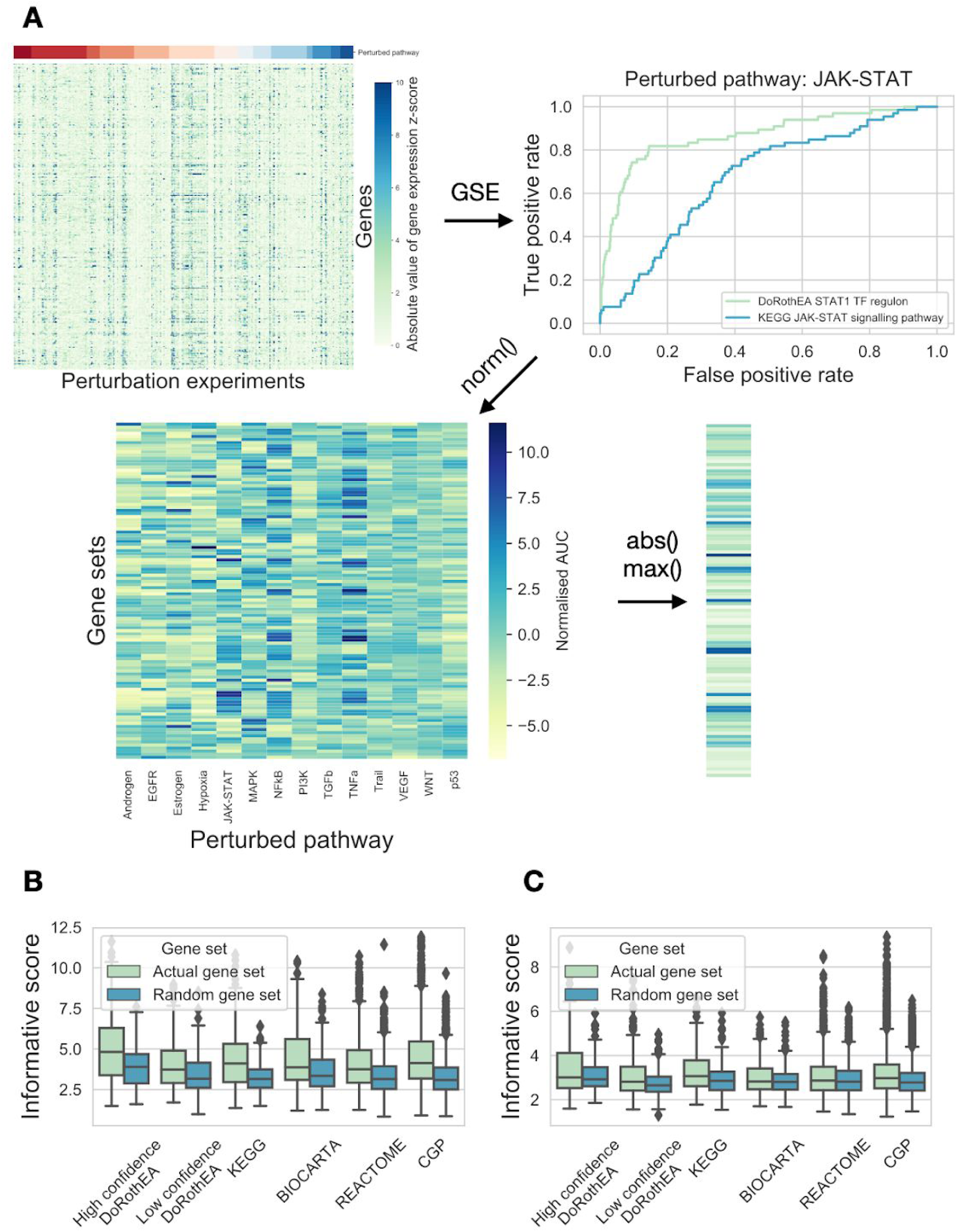
Informative score of gene sets. (A) Schematic pipeline of informative score calculation. From labeled gene expression data (gene expression after pathway perturbation, top left panel), gene set enrichment was performed using different gene sets. The diagnostic ability of gene sets to identify biologically meaningful subgroups (different perturbed pathways) in the data was evaluated with the help of ROC curves (top right left panel). Same analysis was performed for all perturbed pathways - gene set pairs, and normalised based on random ROC curves (bottom left panel). Absolute value was calculated for the gene set - ROC AUC matrix, and for each gene set the maximal value (across biological groups, i.e. perturbed pathways) was given as informative score (bottom right panel). (B and C) Distribution of maximal informative scores (y axis) for gene sets form different gene set databases (x axis) using PROGENy (B) and GDSC (C) benchmark data using absolute gene expression values. Random gene sets (color code) were created by sampling genes from the corresponding gene set database. Analyses with the raw instead of absolute values of the gene expression matrix provided similar results (SFigure 4, STable 5).

### 2.2 Identification of informative gene sets

Once determined the similarities in the components of the gene sets, we studied their information content. We defined informative value of gene sets, based on their ability to identify biologically meaningful differences in gene expression data. For this analysis we required labelled gene expression data according to a biologically defined difference. We used a manually curated perturbation gene expression profile dataset from the PROGENy study (Schubert et al. 2018; Holland, Szalai, and Saez-Rodriguez 2019). In these collected experiments 14 cancer related pathways (EGFR, VEGF, MAPK, PI3K, JAK-STAT, NFkB, TNFa, TGFb, p53, Trail, Hypoxia, Androgen, Estrogen, Wnt) were perturbed (by drugs or genetic perturbations) in a total number of 652 experiments (Figure 3A, top left panel). For each standardised gene expression profile (perturbed vs. control, see Methods) we performed gene set enrichment with the *viper* R package (Alvarez et al. 2016) using the different gene sets of the previous section (2.1). Since standard pathway methods do not use information about the signs of interactions (stimulatory or inhibitory) but the TF-activity methods do, we used absolute values of (standardised) gene expression in both cases so that none has the additional information of the sign.

We then examined the ability of gene sets to identify subgroups (different pathway perturbations) in this gene expression data. We used the normalised enrichment scores (NES) from GSE with the gene set as predicted values, and the known perturbation indicator vector (1 if the pathway was perturbed in the given experiment, 0 otherwise, while gene expression was corrected for the sign of perturbation, see Methods) as true values and performed ROC analysis. As an example, the STAT1 regulon (high confidence DoRothEA) had good performance (Area under the Receiver-Operating Curve (AUROC) = 0.87; Figure 3A, top right panel), while Jak-Stat signaling pathway member genes (KEGG) had a mediocre performance (AUROC = 0.68) for identifying Jak-Stat pathway perturbation experiments. For each gene set, we calculated it’s AUROC for each pathway perturbation experiment (using experiments with the given pathway perturbation as positive samples, Figure 3A, bottom left panel), and normalised it against random AUROC values (see Methods). Finally, we calculated the absolute value of these normalised AUROC values (as low AUROC values, where pathway perturbation is associated with negative enrichment score, is also considered as informative, see Methods). We found that top perturbed pathway - gene set associations are biologically meaningful (STable 2), for example perturbed hypoxia pathway - HIF1A transcription factor (DoRothEA) or perturbed NFkB pathway - Chemokine signaling pathway (KEGG). Finally, to get a single informative score for each gene set, we calculated the maximum value of normalised AUROC values across the perturbed pathways (Figure 3A, right panel). We compared this maximal informative score for all of the used databases (Figure 3B). We found that high confidence DoRothEA gene sets (regulons) had the highest informative score (5.01) and had significant higher informative scores than the other gene set databases (p-values from linear model 5.64e-10, 2.20e-04, 8.20e-09, 3.29e-07 and 1.68e-14 for low confidence DoRothEA, KEGG, Biocarta, Reactome and CGP respectively). In addition, non-random gene sets collectively performed significantly better than random ones (ANOVA p value: 3.65e-258).

As our benchmark dataset (perturbation gene expression profiles) is probably biased to some well studied pathways, we complemented it with an analogous analysis using the gene expression datasets from cancer cell lines in the Genomics of Drug Sensitivity in Cancer (GDSC) study (Iorio et al. 2016). In this case we used AUROC curves to discriminate between cell lines with mutations in key pan-cancer driver genes (total 88 genes, STable 3). The top mutation - gene set associations we obtained were biologically meaningful (STable 4). Calculating the maximal informative scores for the GDSC benchmark data showed similar results than the PROGENy benchmark (Figure 3C). Real gene sets were more informative than random ones (ANOVA p value: 2.86e-58), and high confidence DoRothEA had the highest mean informative score (3.44). In summary, by analysing the informative scores of different gene sets on two benchmark dataset, we found that real gene sets perform better than random ones, and high confidence DoRothEA regulons (footprint based gene sets) had the highest informative score.

### 2.3 Associations between informative score and gene set similarity

After calculating gene set similarities and gene set informative scores, we analysed how these two metrics correspond to each other. To illustrate this analysis, we iteratively selected gene set databases (e.g. high confidence DoRothEA, Figure 4A) and compared the maximal informative score of all other gene sets against their maximal similarity to the selected gene set database. In the case of high confidence DoRothEA, we found that the gene sets similar to high confidence DoRothEA have higher informative scores (Figure 4A, Spearman correlation rho = 0.23, p = 3.71e-65). In contrast, the informative score of genes sets showed negligible correlation with their similarity to Biocarta gene sets (Figure 4B, Spearman correlation rho = 0.03, p = 0.03).

**Figure 4.**
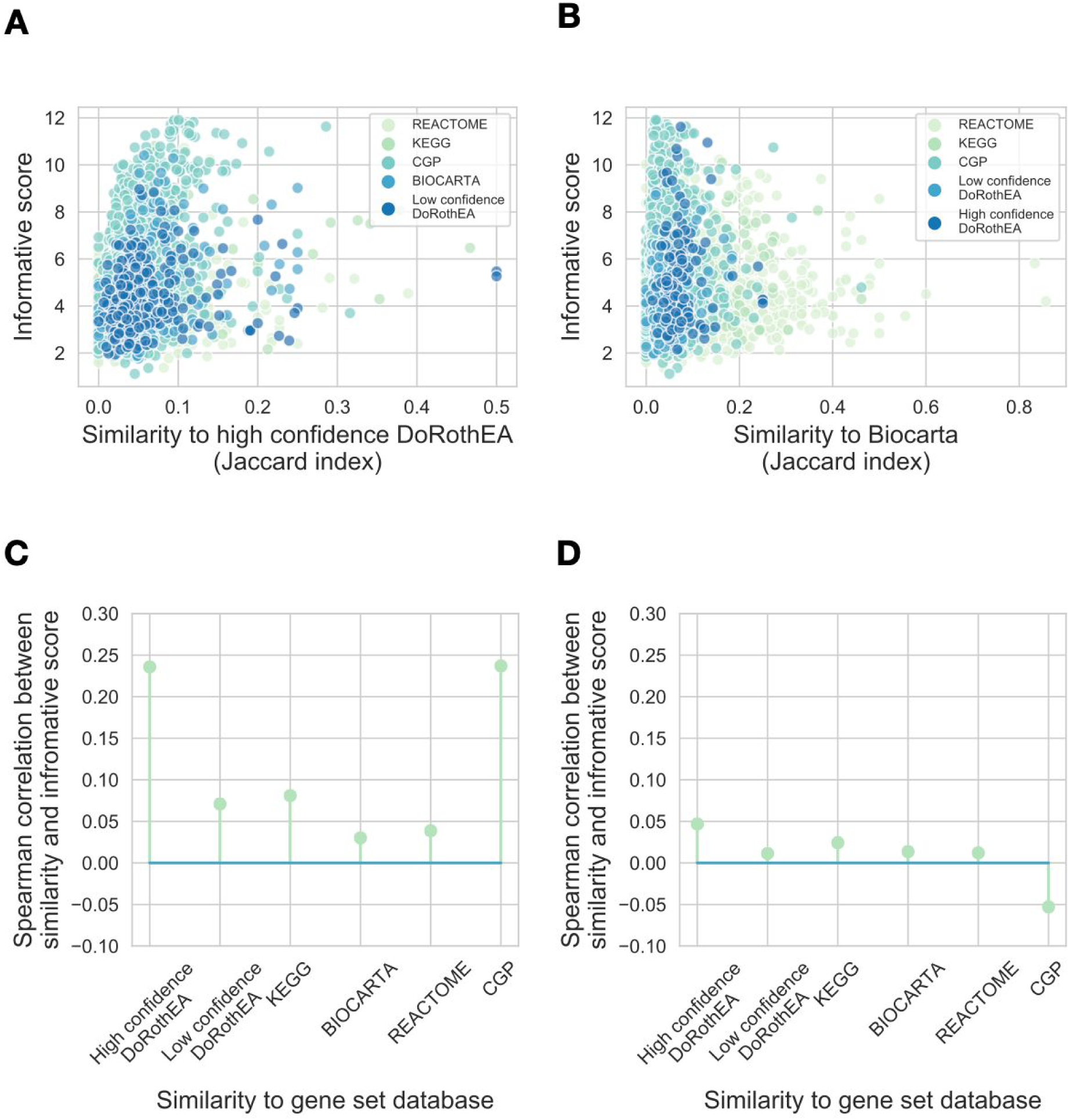
Associations between informative score and gene set similarity. (A and B) Gene set similarity - informative score correlations for high confidence DoRothEA (A) and Biocarta (B). Gene set informative score (y axis) is plotted against similarity (Jaccard index) to high confidence DoRothEA (A) and Biocarta (B) for each gene set from the used gene set databases (color code). (C and D) Spearman correlation coefficients (y axis) between informative score and similarity to gene set database (x axis) are plotted for real gene sets (C) and random gene sets (D). Random gene sets were created by sampling genes from the corresponding gene set database. Analyses with the raw instead of absolute values of the gene expression matrix, using different benchmark datasets (PROGENy or GDSC) and similarity metrics (Jaccard index of overlap coefficient) provided similar results (SFigure 5, STable 6).

We used the PROGENy benchmark dataset for informative score calculation and Jaccard index as similarity metric, and calculated informative score - similarity correlations for each gene set database (Figure 4C). We found that the similarity to high quality DoRothEA and CGP (footprint gene set databases) showed the highest correlation with informative score (Spearman rho = 0.23 and 0.23, Bonferroni corrected p values: 2.23e-64 and 1.65e-29, respectively). In contrast, the similarity to mapping based gene set showed much lower / non-significant correlation with informative score (Spearman correlation rho = 0.08, 0.03 and 0.03, Bonferroni corrected p values: 5.78e-8, 0.2 and 0.16 for KEGG, Biocarta and Reactome, respectively). From these results we concluded that gene sets with high similarity to footprint gene sets have generally higher informative scores. Using randomised gene sets we found no association between gene set similarity and informative score (Figure 4D). In summary, we found that similarity to footprint based but not to mapping based gene sets correlates with the informative score.

### 2.4 Informative scores of gene set differences

As our previous analysis suggested that the informative score of gene sets is correlated with their similarity to footprint based gene sets, we hypothesized that the informative nature of pathway based gene sets is driven by their overlap (ie. similarity) to footprint gene sets. If that is the case, removing the overlapping genes from pathway mapping gene sets is expected to decrease their performance (informative score), but not removing from the footprint gene sets.

To test this hypothesis, we generated set difference gene sets and calculated the informative scores of these new gene sets as described in the previous section (2.2). We also calculated an informative score decrease, a metric showing the decrease of informative score when the overlap with a given gene set was removed (Figure 5A). The set difference gene sets created from high confidence DoRothEA (ie.: the remaining part of high confidence DoRothEA gene sets after removing their intersections with other gene sets) had the highest informative score (Figure 5B, 6.93, linear model p values<1e-100 vs. other gene set databases), followed by set difference gene sets created from CGP (5.58). We did not observe similar behaviour in random gene sets, composed based on the size and genes of the different used gene set databases. Removing the overlap with high confidence DoRothEA regulons resulted in the highest decrease of informative scores of other gene sets (Figure 5C, -0.273 mean informative score decrease, linear model p values <1e-60 vs. other gene set databases), followed by CGP (−0.153 mean informative score decrease), the other footprint based gene set database. We also did not observe similar behaviour in case of random gene sets.

**Figure 5.**
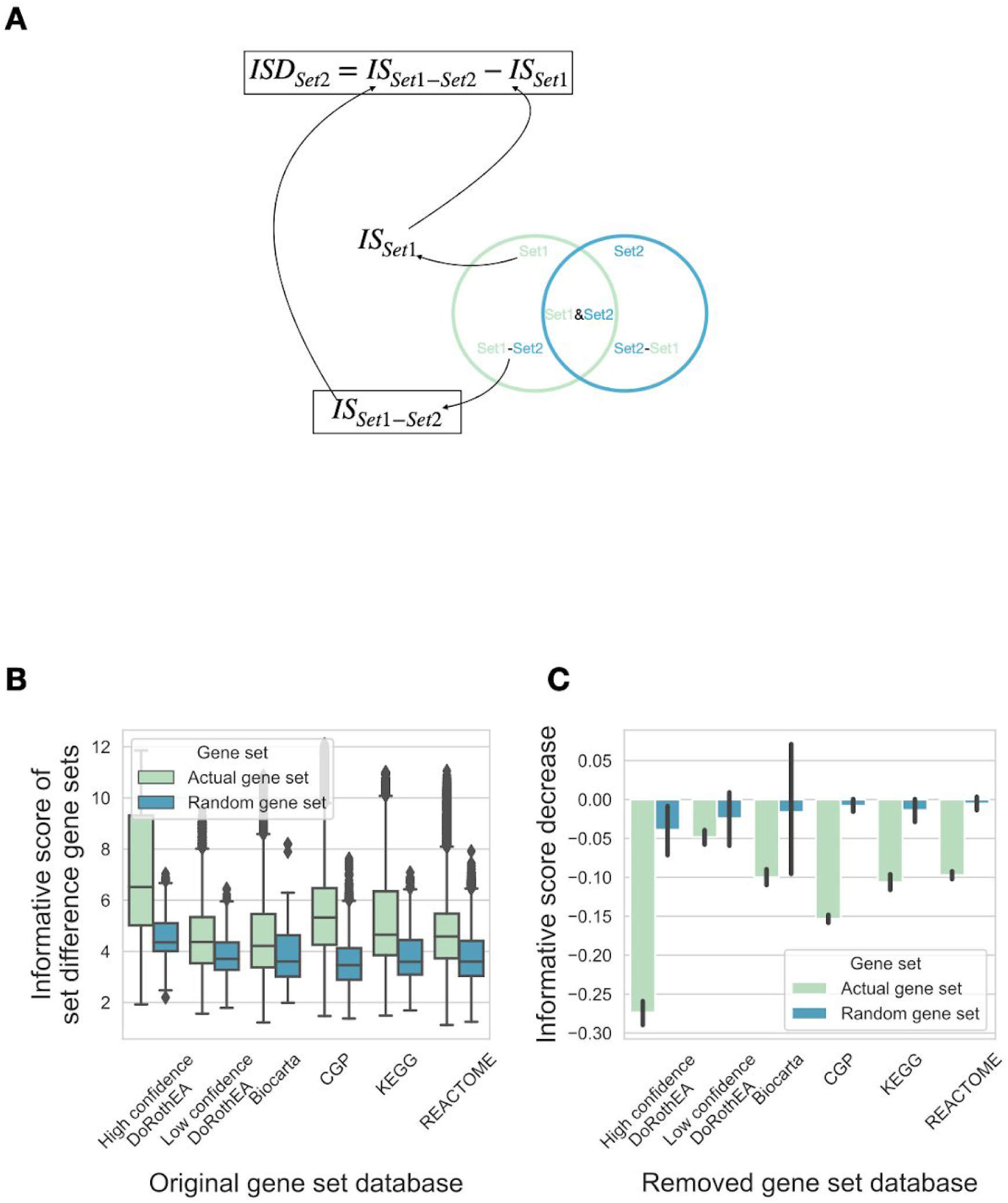
Effect of removing intersection of gene sets. (A) Schematic representation of gene set differences, informative score (IS) and informative score decrease (ISD) calculation. (B) Informative scores of set difference gene sets. For all gene sets from a given database (x axis) set differences with gene sets from other databases were calculated, and informative scores (y axis) were calculated using PROGENy benchmark, absolute gene expression values. Random gene sets (color code) were created by sampling genes from the corresponding gene set database. (C) Informative score decrease was calculated for each gene set from a given gene set database (x axis). Mean informative score +/- 95% CI is shown. Analyses with the raw instead of absolute values of the gene expression matrix and using different benchmark datasets (PROGENy or GDSC) provided similar results (SFigure 6).

Altogether, our results suggest that the main factor driving the informative nature of pathway based gene sets is their overlap with transcription factor regulons / footprint based gene sets.

## 3. Discussion

In this paper, we analysed the overlaps between pathway mapping and footprint based gene sets, and how this overlap influences the usability of gene sets on different benchmark datasets. We found, in concordance with (Cantini et al. 2018), that footprint gene sets are more informative than pathway mapping gene sets (Figure 3). More importantly, we show evidence that the similarity to footprint gene sets influences the informative scores of pathway mapping gene sets (Figure 4). Removing gene intersections with footprint gene sets also significantly decrease the informative value of pathway gene sets (Figure 5). We found that protein products of genes regulated by a given transcription factor also frequently interact in the protein-protein interaction network (Figure 2). The main reason behind this is probably that TFs regulate specific cellular processes, and hence the regulated genes are also interacting with each other on protein level. These results also explain the significant overlap between pathway mapping and footprint based gene sets.

While the effect of different confounding factors (such as gene set size, gene set database size, gene composition of gene sets) can not be fully ruled out, in all of our analysis we used random gene sets (composed from random sampled genes from the original database, with similar gene set size and number of gene sets) as controls. The robustness of our analysis across different benchmarks suggests the importance of differences between footprint and mapping based gene sets.

Of note, several pathway based gene sets contain not only protein-protein interactions, but also gene regulatory interactions. For example KEGG_P53_SIGNALING_PATHWAY (https://www.genome.jp/kegg-bin/show_pathway?hsa04115) contains protein-protein interactions regulating TP53 activity as well as genes regulated by TP53 (TP53 regulon). Not surprisingly, enrichment analysis using this gene set effectively identifies TP53 related changes (TP53 perturbations in PROGENy benchmark and TP53 mutations in GDSC benchmark). Also, we observed a larger effect of similarity to footprint gene sets / removing overlaps with footprint gene sets in our perturbation benchmark (PROGENy) than in the mutation benchmark (GDSC), suggesting that classical pathway based gene sets are better usable with steady state than with perturbation data.

Our analysis suggests that informative pathway mapping based gene sets are not informative based on the biological process (pathway) regulated by the protein product of these genes, but due to their overlap with footprint gene sets, especially transcription factor regulons. These results suggest that different pathway analysis techniques, using gene expression data, might not actually infer pathway activity, but the activity of the overlapping transcription factor, so the results of these methods should be used with caution. For example, gene expression changes can be compensatory (negative feedback) for the original perturbation. Perturbations of the MAPK pathway (e.g.: KRAS / BRAF mutations) frequently lead to overexpression of MAPK phosphatases (DUSPs), and decreases the expression of MAPK pathway members (Schubert et al. 2018). In this case mapping based pathway analysis techniques (especially using methods with network topology) can identify the perturbation of pathway, but can (incorrectly) assume decreased pathway activity.

Protein-protein interaction and other biochemical information give us nevertheless very valuable information to study pathways. We believe that combining gene expression, data driven inference of protein activity and network information (A. Liu et al. 2019; Paull et al. 2013) allows us to extract more accurate insights into signal transduction.

In summary, we found that footprint gene sets perform better than pathway based gene sets in different benchmark data, and that good performance of pathway based gene sets is (partially) explainable based on their overlap with footprint gene sets. Based on these results and on the known weak association between gene expression and protein activity, we submit that gene expression data is better suited for footprint based than mapping-based to estimate the activity of pathways.

## 4. Methods

### Gene sets

We used gene sets (KEGG, BIOCARTA, REACTOME and CGP) from MSigDB and transcription factor regulons from DoRothEA databases. To access MSigDB gene sets we used the *msigdbr* R package. We used high confidence (DoRothEA_AB) and low confidence (DoRothEA_CD) versions of DoRothEA regulon dataset. For each gene set database, we created a randomised control version. The randomised gene set databases contained the same number of gene sets as the original databases, but genes were randomly sampled from the genes of the original database, with random sampled gene set sizes from the original database.

For set difference gene sets we calculated set difference (S_1_ - S_2_) for all pairs of gene sets, where S_1_ and S_2_ were coming from different gene set databases.

### Gene set similarity

We used Jaccard index

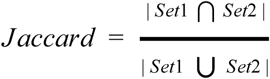

and overlap coefficient

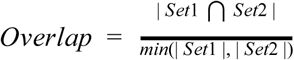

to calculate gene set similarities.

### Shortest path distance calculation

Protein-protein interaction network was downloaded from http://omnipathdb.org/ (Türei, Korcsmáros, and Saez-Rodriguez 2016). OmniPath is a meta-network of literature curated mammalian signaling pathways. Uniprot IDs were converted to gene symbols using the *biomaRt* R package (Durinck et al. 2009). We created an undirected version of the directed signaling network, selected the largest connected component of this network and calculated the shortest path distance between all node pairs. For each gene set (in a given gene set database) we calculated all shortest path distances between the gene set members, and compared it to the whole distance distribution of OmniPath. As genes present in the databases (especially in case of pathway based gene set databases) are expected to have high degree (“well studied” genes), we also compared our observed distance distribution with a degree matched random gene set distribution (as hub genes are expected to have smaller distance with other genes).

### Gene set benchmarking

We defined informative gene sets based on their ability to identify biologically meaningful differences in gene expression data. We used two different gene expression datasets to benchmark the gene sets. We used high quality, manually curated perturbation gene expression data collected for development of the PROGENy method. In this dataset signaling pathways were perturbed, so we used the different pathways as biological subgroups. The other dataset we used was baseline gene expression of cancer cell lines from the GDSC project. In this case we used mutations of several driver genes to create biological subgroups for further analysis.

We calculated normalised enrichment scores (*viper* R package) using the different gene sets and the (standardised) gene expression data. In the case of the GDSC benchmark, we standardised gene expression data across tissue types, to remove tissue type specific expression bias, while in the case of the PROGENy benchmark, we calculated perturbed vs. control gene expression changes. To measure the biological informative score of a gene set, we calculated ROC AUCs. For a given perturbed pathway (PROGENy data) or driver mutation (GDSC data) we selected all occurrences of this pathway or mutation as true positives, and all other samples (were the pathway was not perturbed / mutation was not present) as true negatives for the ROC analysis. We calculated AUROCs for all pathways / mutations and normalised these AUCs to random ROC curves (with the same number of true positive / negative samples) to create ROC AUC z-scores.

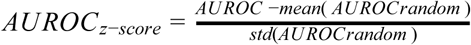

Positive z-scores mean that the expression of gene set members are high in the investigated biological subgroup, while in case of negative z-scores gene set members have low expression. Based on this, gene sets with extreme negative z-scores can also be considered as informative gene sets, so we used the absolute values of these z-scores as final informative scores.

### Informative score - similarity calculation

We selected a gene set database D (eg.: high confidence DoRothEA) and for all other gene sets, we calculated their (maximal) similarity (Jaccard index or overlap coefficient) to D, resulting in a similarity vector. For each gene set, we also calculated their (maximal) informative score (using PROGENy or GDSC benchmark with absolute or raw gene expression values), resulting in an informative score vector. We calculated and reported Spearman rank correlation between similarity vector and informative score vector.

### Informative score decrease calculation

We selected a gene set database D (eg.: high confidence DoRothEA) and for all other gene set databases, we created set difference gene sets S_1-2_ = S_1_ - S_2_, where S_2_ gene sets were coming from D gene set database. We calculated informative score I_1_ for S_1_ and I_1-2_ for S_1-2_, and informative score decrease (for S_2_) as I_1-2_ - I_1_. We reported the mean informative score decrease for D.

## 5. Acknowledgements

BS was supported by the Premium Postdoctoral Fellowship Program of the Hungarian Academy of Sciences [460044]. The authors would like to thank for the usage of MTA Cloud (https://cloud.mta.hu/) that significantly helped us achieve the results published in this paper. We would like to thank Christian Holland and Javier Perales Patón for reading the manuscript and providing useful comments.

## 6. Authors contributions

BS designed the research, performed all analysis and wrote the manuscript. JSR conceptualized and supervised the project and contributed to results interpretation and manuscript writing.

## 7. Code and data availability

Code to reproduce our whole analysis is available at https://github.com/bence-szalai/why-pathway-methods-work.

## Supplementary Materials

**Supplementary Table 1.** Statistics for gene set similarity comparisons. Gene set similarity (Jaccard index or overlap coefficient, 2nd column) values were calculated between DoRothEA gene sets (high and low confidence, 1st column) and other gene sets or random gene sets with corresponding gene size distribution. Mann-Whitney U test was performed between the similarity values for real and random gene sets.

| DoRothEA | Similarity metrics | MWU p value vs. Biocarta | MWU p value vs. KEGG | MWU p value vs. Reactome | MWU p value vs. CGP |
| --- | --- | --- | --- | --- | --- |
| High confidence | Jaccard index | 9.10e-17 | 8.76e-26 | 2.42e-21 | 3.75e-30 |
| High confidence | Overlap coefficient | 5.85e-12 | 9.27e-22 | 2.18e-27 | 1.63e-33 |
| Low confidence | Jaccard index | 4.17e-07 | 7.43e-08 | 3.55e-08 | 9.26e-25 |
| Low confidence | Overlap coefficient | 1.07e-15 | 3.31e-07 | 4.30e-18 | 2.40e-38 |

**Supplementary Table 2.** Top perturbed pathway - gene set associations.

| **High confidence**<br>**DoRothEA** | **Low confidence**<br>**DoRothEA** | **KEGG** | **BIOCARTA** | **REACTOME** | **CGP** |
| --- | --- | --- | --- | --- | --- |
| *Hypoxia HIF1A* | *TNFa RELB* | *Hypoxia*<br>*KEGG_FRUCTOSE_*<br>*AND_MANNOSE_ME*<br>*TABOLISM* | *TNFa*<br>*BIOCARTA_TNFR2_*<br>*PATHWAY* | *Hypoxia*<br>*REACTOME_GLYCO*<br>*LYSIS* | *Hypoxia*<br>*ELVIDGE_HIF1A_TA*<br>*RGETS_DN* |
| *TNFa REL* | *JAK-STAT IRF2* | *Hypoxia*<br>*KEGG_GLYCOLYSIS*<br>*_GLUCONEOGENES*<br>*IS* | *TNFa*<br>*BIOCARTA_CD40_P*<br>*ATHWAY* | *Hypoxia*<br>*REACTOME_GLUCO*<br>*SE_METABOLISM* | *Hypoxia*<br>*FARDIN_HYPOXIA_1*<br>*1* |
| *TNFa RELA* | *JAK-STAT IRF9* | *TNFa*<br>*KEGG_NOD_LIKE_R*<br>*ECEPTOR_SIGNALI*<br>*NG_PATHWAY* | *TNFa*<br>*BIOCARTA_IL1R_PA*<br>*THWAY* | *TNFa*<br>*REACTOME_INTERL*<br>*EUKIN_10_SIGNALI*<br>*NG* | *Hypoxia*<br>*MENSE_HYPOXIA_U*<br>*P* |
| *TNFa NFKB1* | *MAPK HIC1* | *Hypoxia*<br>*KEGG_PENTOSE_P*<br>*HOSPHATE_PATHW*<br>*AY* | *TNFa*<br>*BIOCARTA_NFKB_P*<br>*ATHWAY* | *Hypoxia*<br>*REACTOME_GLYCO*<br>*GEN_SYNTHESIS* | *Hypoxia*<br>*ELVIDGE_HIF1A_AN*<br>*D_HIF2A_TARGETS_*<br>*DN* |
| *JAK-STAT STAT2* | *TNFa NFATC1* | *NFkB*<br>*KEGG_CHEMOKINE*<br>*_SIGNALING_PATH*<br>*WAY* | *TNFa*<br>*BIOCARTA_MSP_PA*<br>*THWAY* | *JAK-STAT*<br>*REACTOME_INTERF*<br>*ERON_ALPHA_BETA*<br>*_SIGNALING* | *Hypoxia*<br>*WACKER_HYPOXIA_*<br>*TARGETS_OF_VHL* |

**Supplementary Table 3.** List of (driver) gene mutations used in GDSC benchmark.

|  |  |  |  |  |  |  |  |  |
| --- | --- | --- | --- | --- | --- | --- | --- | --- |
| ACVR2A | ARID4B | BPTF | CHD9 | FAT1 | MACF1 | MYH9 | PIK3R1 | SPTAN1 |
| AKAP9 | ASH1L | BRAF | CIC | FAT2 | MAGI2 | NCOR1 | PTCH1 | STK11 |
| ALK | ASPM | BRCA1 | CREBBP | FBXW7 | MECOM | NCOR2 | PTEN | TET2 |
| ANK3 | ASXL1 | BRCA2 | CTCF | INPPL1 | MGA | NF1 | RB1 | TP53 |
| APC | ATM | BRWD1 | CTNNB1 | KALRN | MLH1 | NF2 | RNF43 | VHL |
| ARFGEF1 | ATR | CASP8 | DNMT3A | KDM6A | MLL | NOTCH1 | ROBO2 | XRN1 |
| ARHGAP29 | B2M | CDH1 | EGFR | KRAS | MLL2 | NRAS | SACS | ZFHX3 |
| ARID1A | BAP1 | CDKN2A | EP300 | LAMA2 | MLL3 | NSD1 | SETD2 | ZNF292 |
| ARID2 | BCOR | CEP290 | EZH2 | LPHN2 | MLLT4 | PBRM1 | SMAD4 |  |
| ARID4A | BMPR2 | CHD8 | F8 | LRPPRC | MYH11 | PIK3CA | SMARCA4 |  |

**Supplementary Table 4.** Top mutation - gene set associations.

| **High confidence**<br>**DoRothEA** | **Low confidence**<br>**DoRothEA** | **KEGG** | **BIOCARTA** | **REACTOME** | **CGP** |
| --- | --- | --- | --- | --- | --- |
| APC_mut CDX2 | APC_mut RXRB | KRAS_mut<br>KEGG_HISTIDINE_M<br>ETABOLISM | RB1_mut<br>BIOCARTA_GRANUL<br>OCYTES_PATHWAY | KRAS_mut<br>REACTOME_DEFEC<br>TIVE_GALNT3_CAU<br>SES_FAMILIAL_HYP<br>ERPHOSPHATEMIC<br>_TUMORAL_CALCIN<br>OSIS_HFTC | KRAS_mut<br>ANDERSEN_CHOLA<br>NGIOCARCINOMA_<br>CLASS2 |
| BRAF_mut MITF | KRAS_mut SPDEF | RB1_mut<br>KEGG_HEMATOPOI<br>ETIC_CELL_LINEAG<br>E | RB1_mut<br>BIOCARTA_IL1R_PA<br>THWAY | KRAS_mut<br>REACTOME_DEFEC<br>TIVE_C1GALT1C1_C<br>AUSES_TN_POLYAG<br>GLUTINATION_SYN<br>DROME_TNPS | APC_mut<br>WATANABE_COLON<br>_CANCER_MSI_VS_<br>MSS_DN |
| KRAS_mut ERG | APC_mut RXRG | APC_mut<br>KEGG_PHENYLALA<br>NINE_METABOLISM | RB1_mut<br>BIOCARTA_IL5_PAT<br>HWAY | KRAS_mut<br>REACTOME_O_LINK<br>ED_GLYCOSYLATIO<br>N_OF_MUCINS | KRAS_mut<br>VECCHI_GASTRIC_<br>CANCER_ADVANCE<br>D_VS_EARLY_DN |
| BRAF_mut SOX10 | KRAS_mut EHF | RB1_mut<br>KEGG_LONG_TERM<br>_DEPRESSION | KRAS_mut<br>BIOCARTA_NUCLEA<br>RRS_PATHWAY | RB1_mut<br>REACTOME_POTAS<br>SIUM_CHANNELS | APC_mut<br>SABATES_COLORE<br>CTAL_ADENOMA_U<br>P |
| KRAS_mut CDX2 | KRAS_mut ELF3 | RB1_mut<br>KEGG_TASTE_TRAN<br>SDUCTION | RB1_mut<br>BIOCARTA_INFLAM<br>_PATHWAY | KRAS_mut<br>REACTOME_TERMI<br>NATION_OF_O_GLY<br>CAN_BIOSYNTHESI<br>S | KRAS_mut<br>HINATA_NFKB_TAR<br>GETS_KERATINOCY<br>TE_DN |

**Supplementary Table 5.** Statistical comparison of gene set informative scores. Linear model (informative score ∼ gene set database + random) was fitted on informative scores from different benchmark data (rows). ANOVA p values (log10 transformed), linear model coefficients and linear model p values (log10 transformed) are reported. For the gene set database factor, high confidence DoRothEA was the reference level.

|  | ANOVA log10 p value |  | Linear model coefficient |  |  |  |  |  | Linear model log10 p values |  |  |  |  |
| --- | --- | --- | --- | --- | --- | --- | --- | --- | --- | --- | --- | --- | --- |
|  | Random vs.<br>real | Gene sets | DoRothEA<br>High conf. | DoRothEA<br>Low conf. | KEGG | BiOCARTA | REACTOME | CGP | DoRothEA<br>Low conf. | KEGG | BiOCARTA | REACTOME | CGP |
| PROGENy<br>absolute | -257.44 | -18.04 | 5.01 | -0.75 | -0.44 | -0.59 | -0.65 | -0.8 | -9.25 | -3.66 | -8.09 | -6.48 | -13.77 |
| PREOGNy | -129.73 | -10.69 | 5.8 | -0.42 | -0.37 | -0.5 | -0.49 | -0.7 | -2.59 | -2.21 | -4.63 | -3.1 | -8.28 |
| GDSC<br>absolute | -57.54 | -7.68 | 3.44 | -0.39 | -0.36 | -0.25 | -0.18 | -0.29 | -7.37 | -6.75 | -4.46 | -1.84 | -5.58 |
| GDSC | -4.7 | -33.96 | 2.34 | -0.01 | 0.16 | 0.22 | 0.19 | 0.29 | -0.12 | -3.58 | -7.88 | -4.07 | -12.59 |

**Supplementary Table 6.**
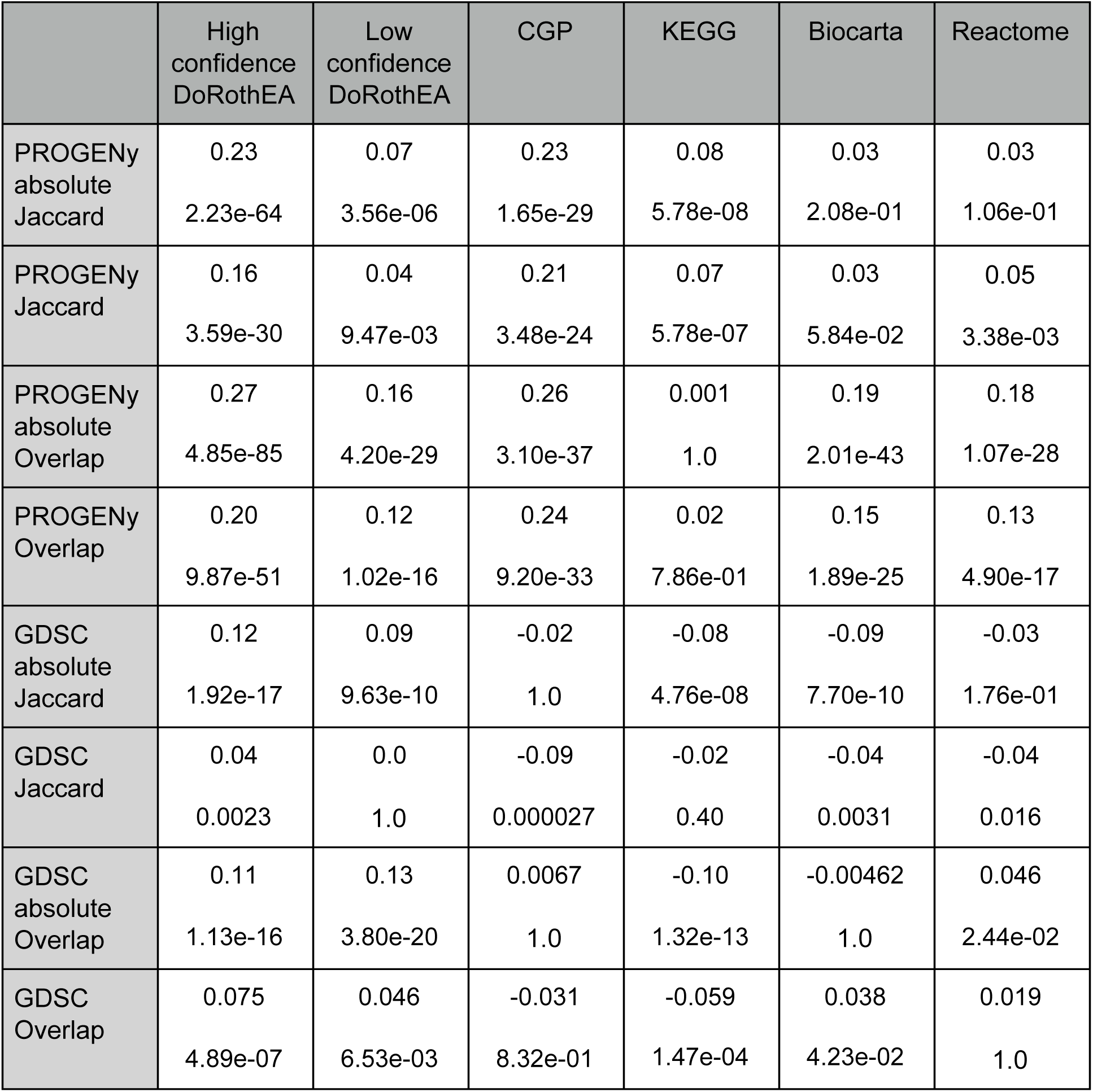
Associations between informative score and gene set similarity. Spearman correlation coefficients (first value) and Bonferroni corrected p values (second value) were calculated between informative scores and similarity to gene set databases (columns). Different benchmark data (PROGENy or GDSC), gene expression data preprocessing (absolute or row value) and similarity metrics (Jaccard index or overlap coefficient) was used (rows).

|  | High confidence<br>DoRothEA | Low confidence<br>DoRothEA | CGP | KEGG | Biocarta | Reactome |
| --- | --- | --- | --- | --- | --- | --- |
| PROGENy<br>absolute<br>Jaccard | 0.23<br>2.23e-64 | 0.07<br>3.56e-06 | 0.23<br>1.65e-29 | 0.08<br>5.78e-08 | 0.03<br>2.08e-01 | 0.03<br>1.06e-01 |
| PROGENy<br>Jaccard | 0.16<br>3.59e-30 | 0.04<br>9.47e-03 | 0.21<br>3.48e-24 | 0.07<br>5.78e-07 | 0.03<br>5.84e-02 | 0.05<br>3.38e-03 |
| PROGENy<br>absolute<br>Overlap | 0.27<br>4.85e-85 | 0.16<br>4.20e-29 | 0.26<br>3.10e-37 | 0.001<br>1.0 | 0.19<br>2.01e-43 | 0.18<br>1.07e-28 |
| PROGENy<br>Overlap | 0.20<br>9.87e-51 | 0.12<br>1.02e-16 | 0.24<br>9.20e-33 | 0.02<br>7.86e-01 | 0.15<br>1.89e-25 | 0.13<br>4.90e-17 |
| GDSC<br>absolute<br>Jaccard | 0.12<br>1.92e-17 | 0.09<br>9.63e-10 | -0.02<br>1.0 | -0.08<br>4.76e-08 | -0.09<br>7.70e-10 | -0.03<br>1.76e-01 |
| GDSC<br>Jaccard | 0.04<br>0.0023 | 0.0<br>1.0 | -0.09<br>0.000027 | -0.02<br>0.40 | -0.04<br>0.0031 | -0.04<br>0.016 |
| GDSC<br>absolute<br>Overlap | 0.11<br>1.13e-16 | 0.13<br>3.80e-20 | 0.0067<br>1.0 | -0.10<br>1.32e-13 | -0.00462<br>1.0 | 0.046<br>2.44e-02 |
| GDSC<br>Overlap | 0.075<br>4.89e-07 | 0.046<br>6.53e-03 | -0.031<br>8.32e-01 | -0.059<br>1.47e-04 | 0.038<br>4.23e-02 | 0.019<br>1.0 |

**Supplementary Figure 1.**
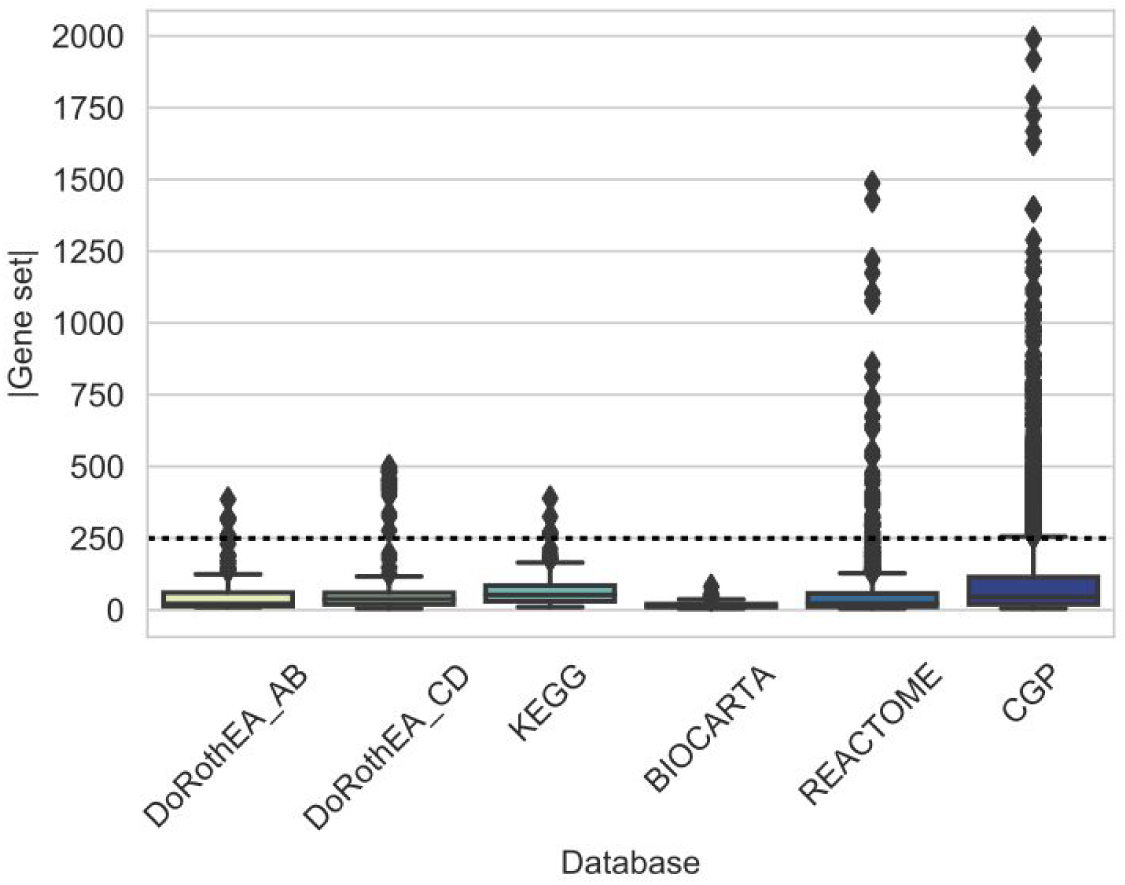
Size of gene sets in the different used gene set databases.

**Supplementary Figure 2.**
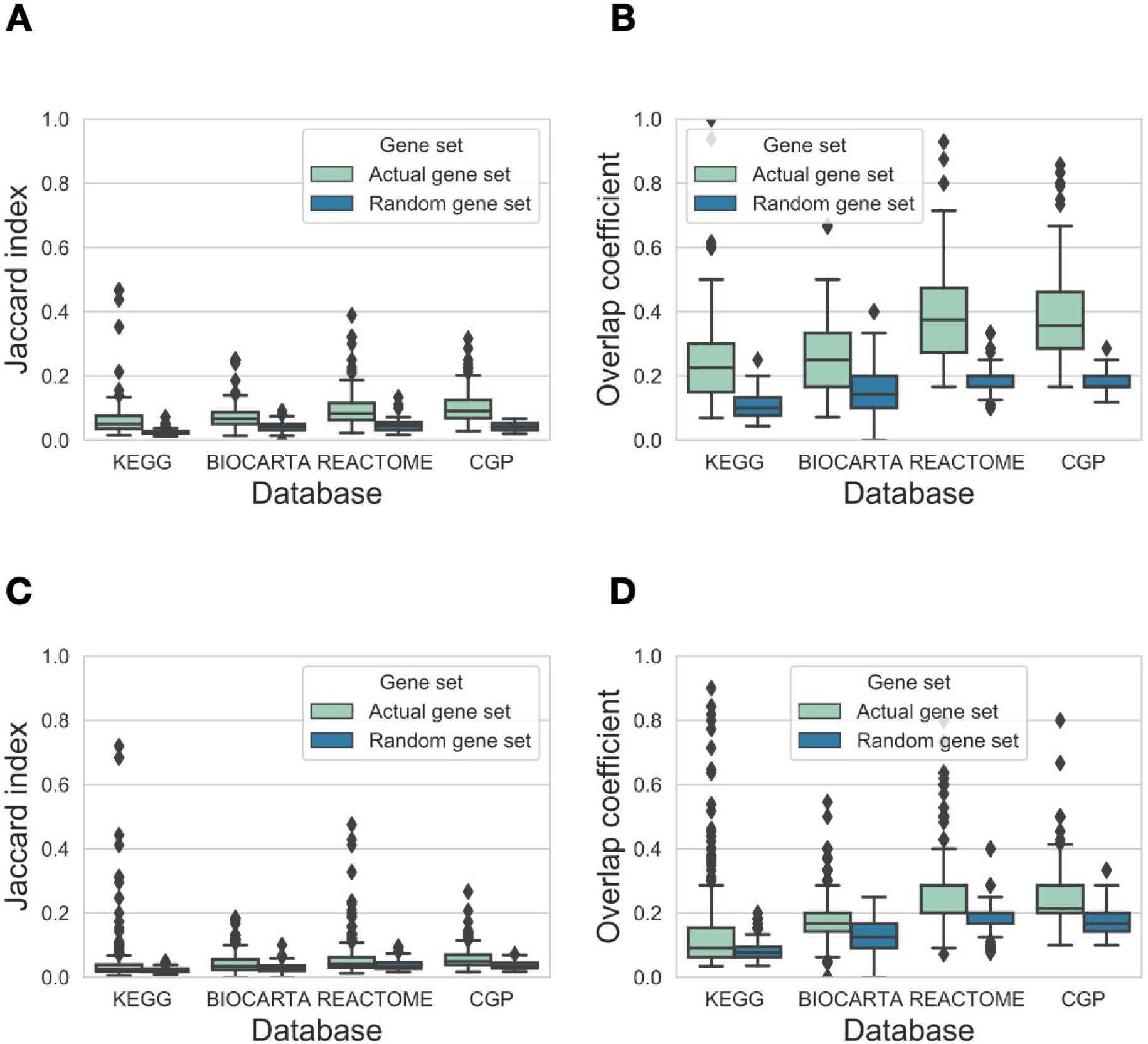
Similarity between DoRothEA regulons and other gene set resources. Distribution of maximal similarity values (y axis, Jaccard index for A and C, overlap coefficient for B and D) to high confidence (A, B) or low confidence (C, D) DoRothEA regulons for the investigated gene set databases (x axis). Random gene sets (color code) were created by sampling gene sets (with corresponding size) from the genes of the given database.

**Supplementary Figure 3.**
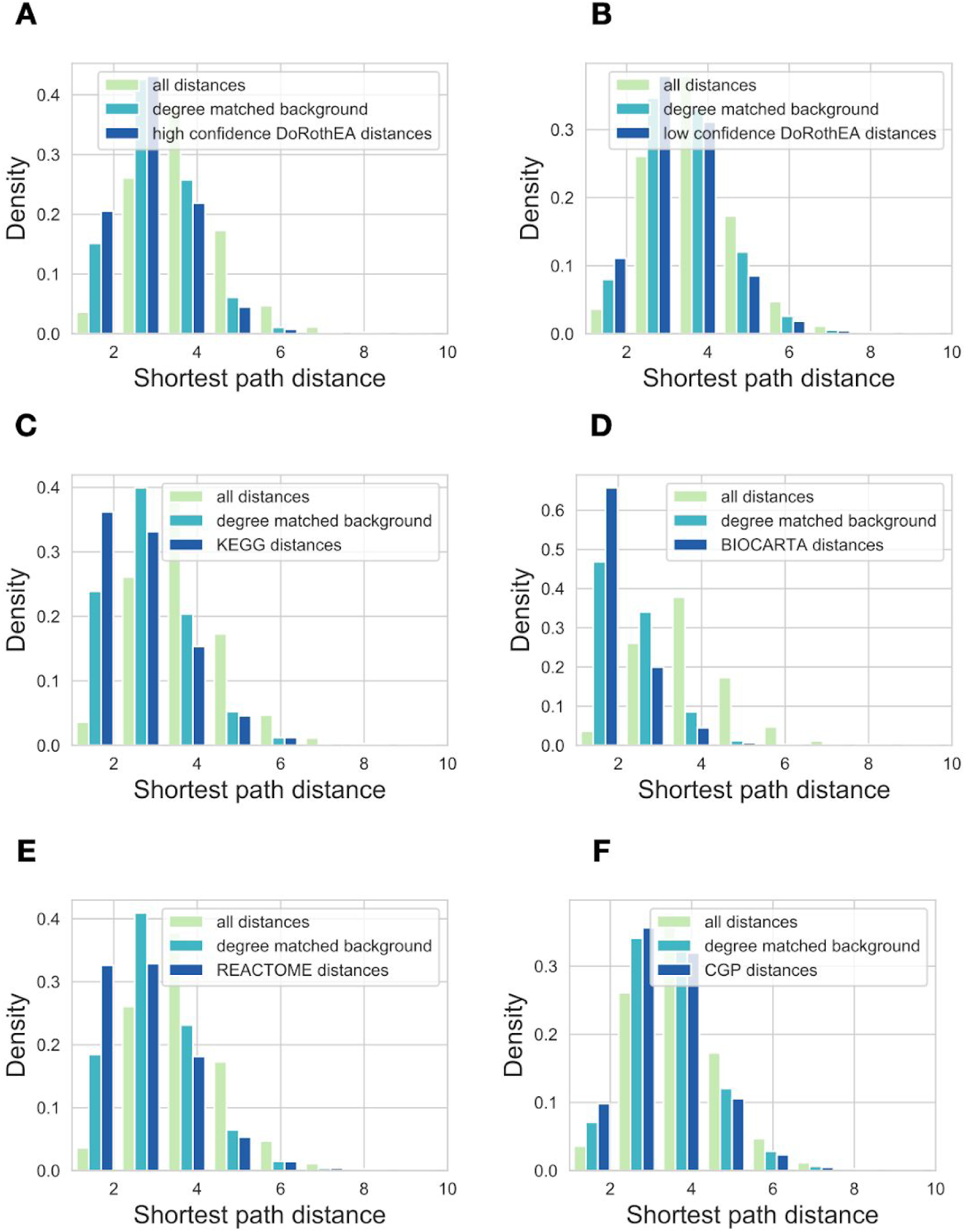
Shortest path distances for gene sets members. Distribution of the shortest path distances for proteins from the same gene set. Used gene set databases: (A) high confidence DoRothEA (B) low confidence DoRothEA (C) KEGG pathways (D) Biocarta (E) Reactome (F) CGP.

**Supplementary Figure 4.**
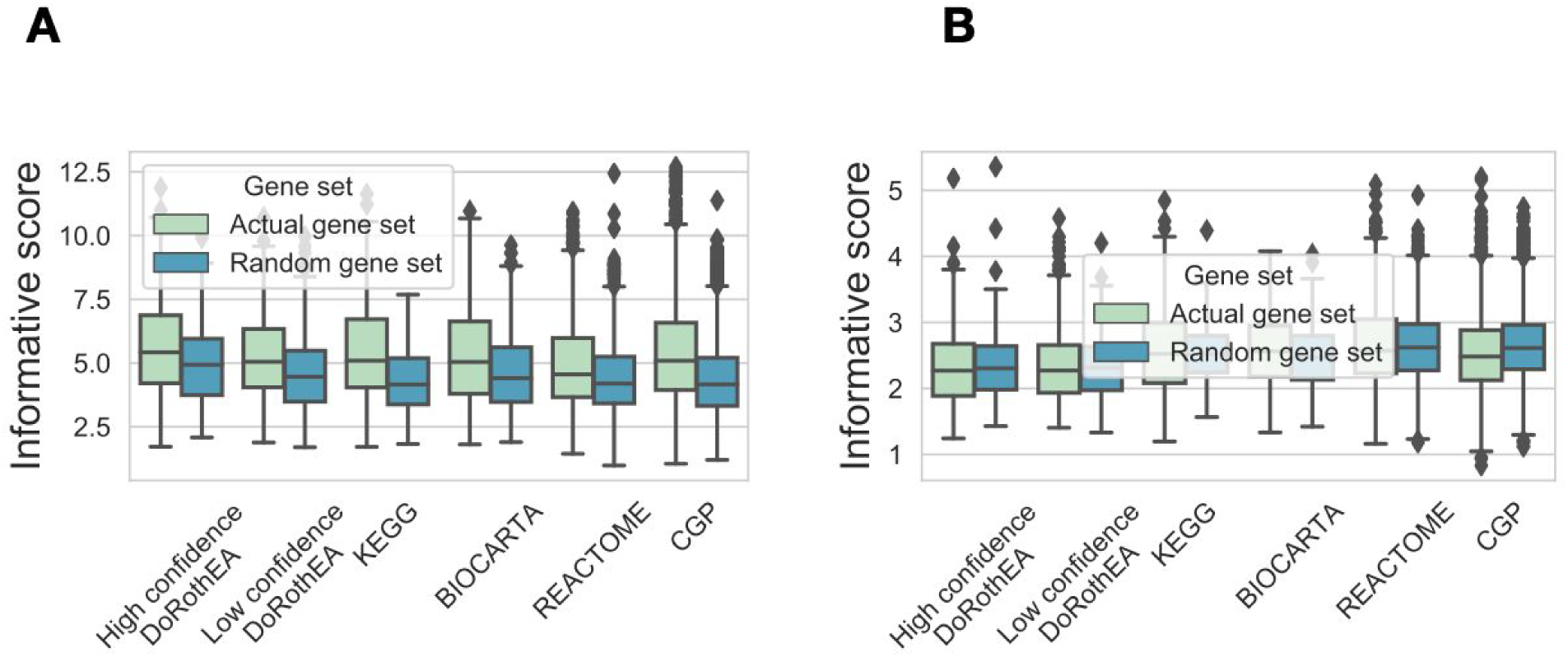
Distribution of maximal informative scores. Distribution of maximal informative scores (y axis) for gene sets form different gene set databases (x axis) using PROGENy (B) and GDSC (C) benchmark data using raw gene expression values. Random gene sets (color code) were created by sampling genes from the corresponding gene set database.

**Supplementary Figure 5.**
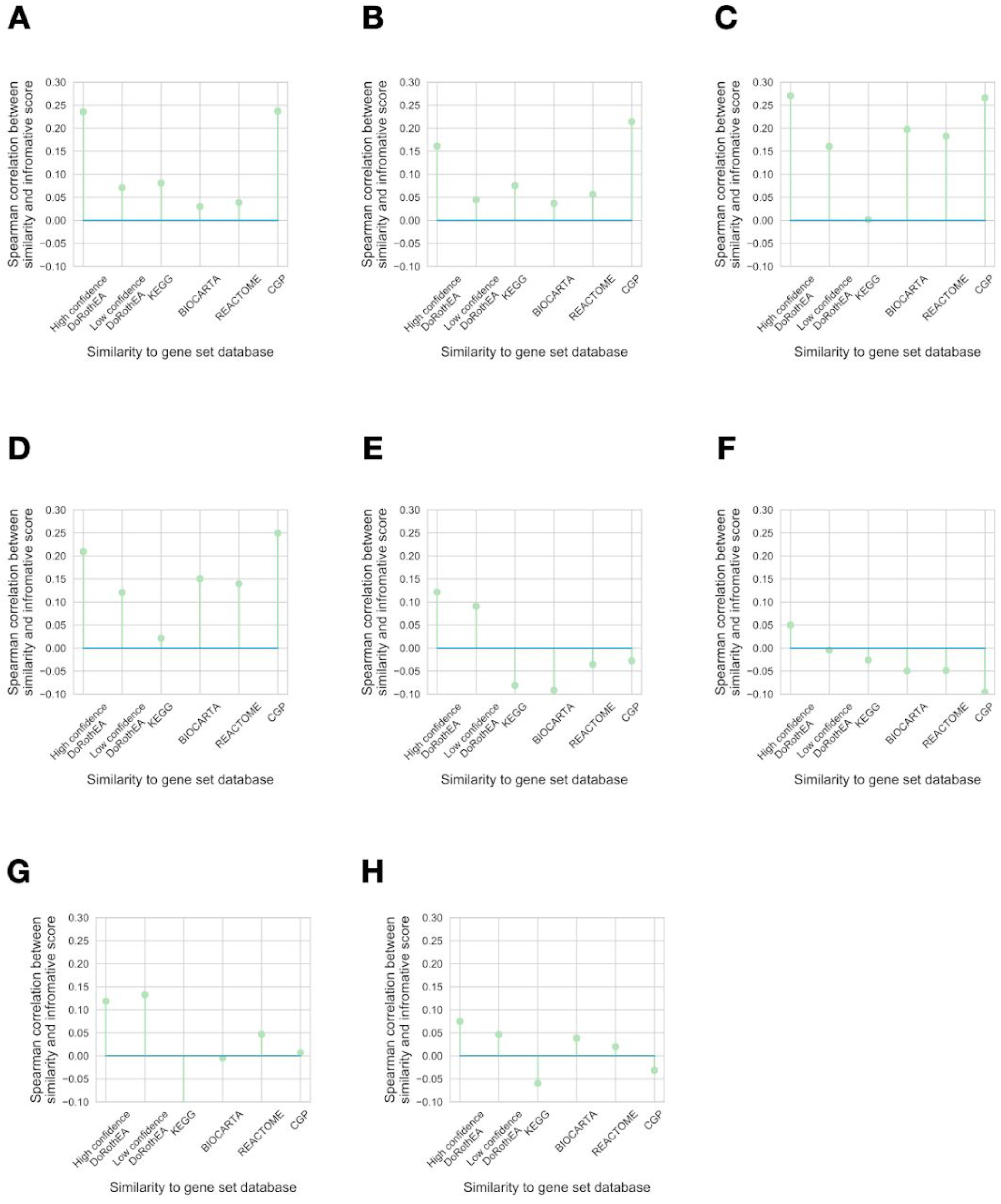
Associations between informative score and gene set similarity. Spearman correlation coefficients (y axis) between informative score and similarity to gene set database (x axis) are plotted for using PROGENy (A, B, C, D) or GDSC (E, F, G, H) benchmark data, absolute (A, C, E, G) or raw (B, D, F, H) gene expression values and Jaccard index (A, B, E, F) or overlap coefficient (C, D, G, H) as similarity metric.

**Supplementary Figure 6.**
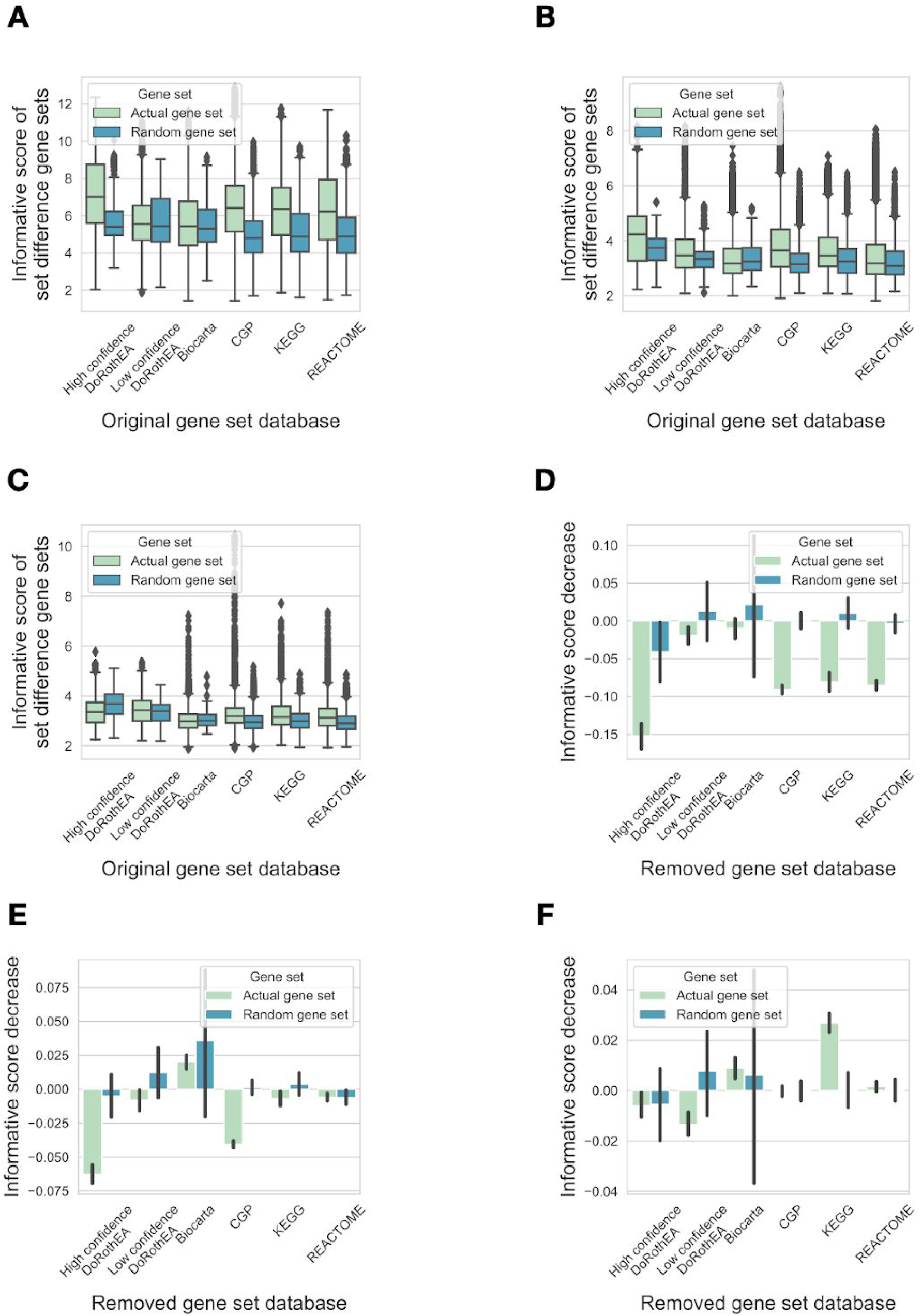
Effect of removing intersection of gene sets. Informative scores of set difference gene sets (A, B, C). For all gene sets from a given database (x axis) set differences with gene sets from other databases were calculated, and informative scores (y axis) were calculated using PROGENy (A) or GDSC benchmark (B, C), and raw (A, C) or absolute (B) gene expression values. Random gene sets (color code) were created by sampling genes from the corresponding gene set database. (C) Informative score decrease was calculated for each gene set from a given gene set database (x axis). Mean informative score +/- 95% CI is shown. PROGENy (D) or GDSC (E, F) benchmark and raw (D, F) of absolute (E) gene expression values were used for informative score calculation.

## Notes

### Competing Interest Statement

The authors have declared no competing interest.

## References

Alvarez, Mariano J., Yao Shen, Federico M. Giorgi, Alexander Lachmann, B. Belinda Ding, B. Hilda Ye, and Andrea Califano. 2016. “Functional Characterization of Somatic Mutations in Cancer Using Network-Based Inference of Protein Activity.” Nature Genetics 48 (8): 838–47.

Buccitelli, Christopher, and Matthias Selbach. 2020. “mRNAs, Proteins and the Emerging Principles of Gene Expression Control.” Nature Reviews. Genetics, July. https://doi.org/10.1038/s41576-020-0258-4.

Cantini, Laura, Laurence Calzone, Loredana Martignetti, Mattias Rydenfelt, Nils Blüthgen, Emmanuel Barillot, and Andrei Zinovyev. 2018. “Classification of Gene Signatures for Their Information Value and Functional Redundancy.” NPJ Systems Biology and Applications 4: 2.

Dugourd, A., and J. Saez-Rodriguez. 2019. “Footprint-Based Functional Analysis of Multi-Omic Data.” Current Opinion in Systems Biology. https://www.sciencedirect.com/science/article/pii/S2452310019300149.

Durinck, Steffen, Paul T. Spellman, Ewan Birney, and Wolfgang Huber. 2009. “Mapping Identifiers for the Integration of Genomic Datasets with the R/Bioconductor Package biomaRt.” Nature Protocols 4 (8): 1184–91.

Garcia-Alonso, Luz, Christian H. Holland, Mahmoud M. Ibrahim, Denes Turei, and Julio Saez-Rodriguez. 2019. “Benchmark and Integration of Resources for the Estimation of Human Transcription Factor Activities.” Genome Research 29 (8): 1363–75.

Garcia-Alonso, Luz, Francesco Iorio, Angela Matchan, Nuno Fonseca, Patricia Jaaks, Gareth Peat, Miguel Pignatelli, et al. 2018. “Transcription Factor Activities Enhance Markers of Drug Sensitivity in Cancer.” Cancer Research 78 (3): 769–80.

Hidalgo, Marta R., Cankut Cubuk, Alicia Amadoz, Francisco Salavert, José Carbonell-Caballero, and Joaquin Dopazo. 2017. “High Throughput Estimation of Functional Cell Activities Reveals Disease Mechanisms and Predicts Relevant Clinical Outcomes.” Oncotarget 8 (3): 5160–78.

Holland, Christian H., Bence Szalai, and Julio Saez-Rodriguez. 2019. “Transfer of Regulatory Knowledge from Human to Mouse for Functional Genomics Analysis.” Biochimica et Biophysica Acta, Gene Regulatory Mechanisms, September, 194431.

Holland, Christian H., Jovan Tanevski, Javier Perales-Patón, Jan Gleixner, Manu P. Kumar, Elisabetta Mereu, Brian A. Joughin, et al. 2020. “Robustness and Applicability of Transcription Factor and Pathway Analysis Tools on Single-Cell RNA-Seq Data.” Genome Biology 21 (1): 36.

Iorio, Francesco, Theo A. Knijnenburg, Daniel J. Vis, Graham R. Bignell, Michael P. Menden, Michael Schubert, Nanne Aben, et al. 2016. “A Landscape of Pharmacogenomic Interactions in Cancer.” Cell 166 (3): 740–54.

Jassal, Bijay, Lisa Matthews, Guilherme Viteri, Chuqiao Gong, Pascual Lorente, Antonio Fabregat, Konstantinos Sidiropoulos, et al. 2020. “The Reactome Pathway Knowledgebase.” Nucleic Acids Research 48 (D1): D498–503.

Kanehisa, Minoru, Yoko Sato, Miho Furumichi, Kanae Morishima, and Mao Tanabe. 2019. “New Approach for Understanding Genome Variations in KEGG.” Nucleic Acids Research 47 (D1): D590–95.

Keenan, Alexandra B., Denis Torre, Alexander Lachmann, Ariel K. Leong, Megan L. Wojciechowicz, Vivian Utti, Kathleen M. Jagodnik, Eryk Kropiwnicki, Zichen Wang, and Avi Ma’ayan. 2019. “ChEA3: Transcription Factor Enrichment Analysis by Orthogonal Omics Integration.” Nucleic Acids Research 47 (W1): W212–24.

Larsen, Simon J., Richard Röttger, Harald H. H. W. Schmidt, and Jan Baumbach. 2019. “E. Coli Gene Regulatory Networks Are Inconsistent with Gene Expression Data.” Nucleic Acids Research 47 (1): 85–92.

Liu, Anika, Panuwat Trairatphisan, Enio Gjerga, Athanasios Didangelos, Jonathan Barratt, and Julio Saez-Rodriguez. 2019. “From Expression Footprints to Causal Pathways: Contextualizing Large Signaling Networks with CARNIVAL.” NPJ Systems Biology and Applications 5 (November): 40.

Liu, Yansheng, Andreas Beyer, and Ruedi Aebersold. 2016. “On the Dependency of Cellular Protein Levels on mRNA Abundance.” Cell 165 (3): 535–50.

Nguyen, Tuan-Minh, Adib Shafi, Tin Nguyen, and Sorin Draghici. 2019. “Identifying Significantly Impacted Pathways: A Comprehensive Review and Assessment.” Genome Biology 20 (1): 203.

Nusinow, David P., John Szpyt, Mahmoud Ghandi, Christopher M. Rose, E. Robert McDonald 3rd, Marian Kalocsay, Judit Jané-Valbuena, et al. 2020. “Quantitative Proteomics of the Cancer Cell Line Encyclopedia.” Cell 180 (2): 387–402.e16.

Parikh, Jignesh R., Bertram Klinger, Yu Xia, Jarrod A. Marto, and Nils Blüthgen. 2010. “Discovering Causal Signaling Pathways through Gene-Expression Patterns.” Nucleic Acids Research 38 (Web Server issue): W109–17.

Paull, Evan O., Daniel E. Carlin, Mario Niepel, Peter K. Sorger, David Haussler, and Joshua M. Stuart. 2013. “Discovering Causal Pathways Linking Genomic Events to Transcriptional States Using Tied Diffusion Through Interacting Events (TieDIE).” Bioinformatics 29 (21): 2757–64.

Piran, Mehran, Reza Karbalaei, Mehrdad Piran, Jehad Aldahdooh, Mehdi Mirzaie, Naser Ansari-Pour, Jing Tang, and Mohieddin Jafari. 2020. “Can We Assume the Gene Expression Profile as a Proxy for Signaling Network Activity?” Biomolecules 10 (6). https://doi.org/10.3390/biom10060850.

Saez-Rodriguez, Julio, and Nils Blüthgen. 2020. “Personalized Signaling Models for Personalized Treatments.” Molecular Systems Biology 16 (1): e9042.

Schubert, Michael, Bertram Klinger, Martina Klünemann, Anja Sieber, Florian Uhlitz, Sascha Sauer, Mathew J. Garnett, Nils Blüthgen, and Julio Saez-Rodriguez. 2018. “Perturbation-Response Genes Reveal Signaling Footprints in Cancer Gene Expression.” Nature Communications 9 (1): 20.

Smith, Joan C., and Jason M. Sheltzer. 2018. “Systematic Identification of Mutations and Copy Number Alterations Associated with Cancer Patient Prognosis.” eLife 7 (December). https://doi.org/10.7554/eLife.39217.

Subramanian, Aravind, Pablo Tamayo, Vamsi K. Mootha, Sayan Mukherjee, Benjamin L. Ebert, Michael A. Gillette, Amanda Paulovich, et al. 2005. “Gene Set Enrichment Analysis: A Knowledge-Based Approach for Interpreting Genome-Wide Expression Profiles.” Proceedings of the National Academy of Sciences of the United States of America 102 (43): 15545–50.

Türei, Dénes, Tamás Korcsmáros, and Julio Saez-Rodriguez. 2016. “OmniPath: Guidelines and Gateway for Literature-Curated Signaling Pathway Resources.” Nature Methods 13 (12): 966–67.

Yaffe, Michael B. 2019. “Why Geneticists Stole Cancer Research Even Though Cancer Is Primarily a Signaling Disease.” Science Signaling 12 (565). https://doi.org/10.1126/scisignal.aaw3483.

Yang, Mi, Francesca Petralia, Zhi Li, Hongyang Li, Weiping Ma, Xiaoyu Song, Sunkyu Kim, et al. 2020. “Crowdsourced Assessment of the of Predictability of Cancer Protein and Phosphoprotein Levels from Genomics and Transcriptomics.” Cell Systems, July. https://doi.org/10.1016/j.cels.2020.06.013.

